# A CDC-42-regulated actin network is necessary for nuclear migration through constricted spaces in *C. elegans*

**DOI:** 10.1101/2023.06.22.546138

**Authors:** Jamie Ho, Leslie A. Guerrero, Diana Libuda, GW Gant Luxton, Daniel A Starr

## Abstract

Successful nuclear migration through constricted spaces between cells or in the extracellular matrix relies on the ability of the nucleus to deform. Little is known of how this takes place *in vivo*. We study confined nuclear migration in *Caenorhabditis elegans* larval P-cells, which is mediated by the LINC complex to pull nuclei towards the minus ends of microtubules. Null mutations of LINC component *unc-84* lead to a temperature-dependent phenotype, suggesting a parallel pathway for P-cell nuclear migration. A forward genetic screen for enhancers of *unc-84* identified *cgef-1* (**C**DC-42 **G**uanine Nucleotide **E**xchange **F**actor). Knockdown of CDC-42 in the absence of the LINC complex led to a P-cell nuclear migration defect. Expression of constitutively active CDC-42 rescued nuclear migration in *cgef-1; unc-84* double mutants suggesting CDC-42 functions downstream of CGEF-1. The Arp2/3 complex and non-muscle myosin II (NMY-2) were also found to function parallel to the LINC pathway. In our model, CGEF-1 activates CDC-42, induces actin polymerization through the Arp2/3 complex to deform the nucleus during nuclear migration while NMY-2 helps push the nucleus through confined spaces.

## Introduction

Cellular migration through constricted spaces is a process that occurs during the immune response, tissue development, and cancer metastasis (Bone & Starr, 2016; Denais et al., 2016; Thiam et al., 2016). During mammalian brain development, newly born neurons must migrate from the germinal layers to the developing cortices by migrating through constrictions generated by the surrounding neural tissue (Kalukula et al., 2022; Kengaku, 2018). Additionally, during the immune response, neutrophils in the blood stream must migrate through the endothelial monolayer of the blood vessels in order to access the site of inflammation or tissue injury (Liu et al., 2021; Salvermoser et al., 2018). The rate of cellular migration through narrow spaces is limited by nuclear deformability as the nucleus is the largest and most rigid organelle of the cell (Friedl et al., 2011; Fu et al., 2012; Swift et al., 2013; Wolf et al., 2013). Nuclear deformability is dependent on several factors such as lamin composition, levels of heterochromatin, and cytoskeletal dynamics. Low expression or knockdown of laminA/C or low levels of heterochromatin results in increased nuclear deformability (Bell et al., 2022; Davidson et al., 2014, 2015; Stephens et al., 2018). Additionally, cytoskeletal forces applied to nuclei can affect nuclear deformability (Renkawitz et al., 2019; Thiam et al., 2016). Mouse dendritic cells that are induced to migrate through constrictions are unable to undergo successful nuclear migration when the actin-nucleating Arp2/3 complex is inhibited (Thiam et al., 2016), highlighting the importance of actin during this process. In addition, nuclear migration through constricted spaces lead to nuclear envelope rupture and increased DNA damage (Denais et al., 2016; Raab et al., 2016; Thiam et al., 2016). Most of these findings were made using *in vitro* systems of cells migrating through manufactured constricted spaces. A system where mouse dendritic cells are imaged migrating through the extra-cellular matrix of explanted mouse ears has been used to find more *in vivo* relevance (Raab et al., 2016). However, how cells and nuclei migrate through constricted spaces as a normal part of development *in vivo* is poorly understood.

We developed an *in vivo* model to study nuclear migration through constricted spaces using larval hypodermal precursor cells (P cells) in *C. elegans* (Fig. 1A)(Bone et al., 2016; Chang et al., 2013). During the early L1 larval stage, twelve P cells organized into six pairs span the lateral side to the ventral side of the animal, with the nuclei located on the lateral side of the animal (Sulston & Horvitz, 1977). During the mid-L1 development, P-cell nuclei, which are 3-4 μm in diameter, migrate from their lateral positions to the ventral cord by squeezing through a narrow space of ∼200 nm between the body wall muscles and the cuticle (Bone et al., 2016; Cox & Hardin, 2004; Francis & Waterston, 1991). This constriction is about 5% the diameter of the nucleus. After P-cell nuclei successfully migrate, they divide and develop into vulval cells and GABA neurons. Failed nuclear migration results in P-cell death, which results in the lack of a vulva and GABA neurons, leading to **egg l**aying deficient (Egl) and **unc**oordinated (Unc) phenotypes (Horvitz & Sulston, 1980; Sulston & Horvitz, 1981).

**Figure 1.**
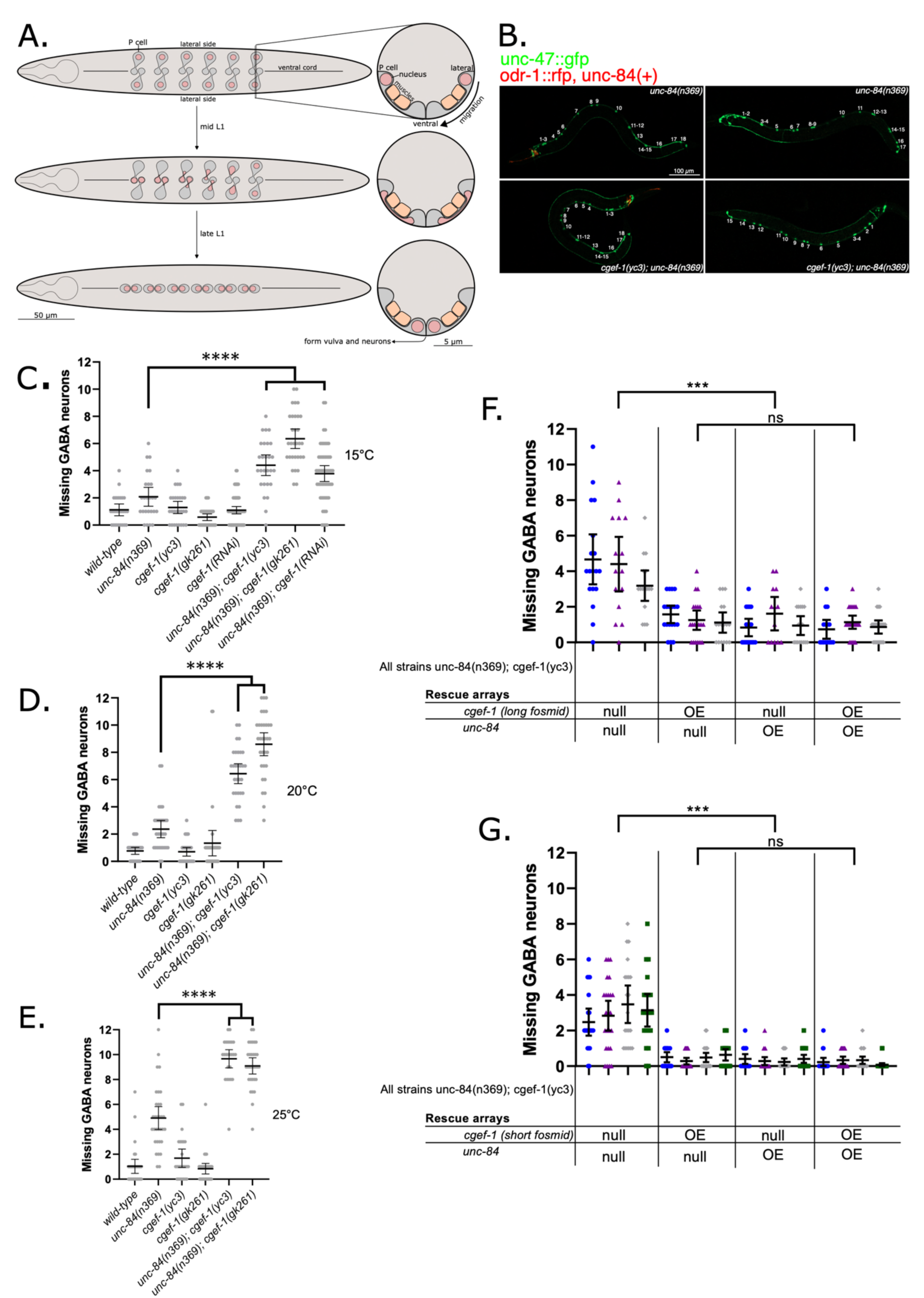
Mutations in *cgef-1* enhance the P-cell nuclear migration defect of *unc-84(null)*. **(A).** Schematic of P-cell nuclear migration. On the left are ventral views of L1 larva during P-cell nuclear migration. Anterior is shown on the left where a pharynx is drawn. The ventral cord is marked with a line down the center of the larvae. Shown on the right are cross sections of analogous stages with the ventral surface of the animal facing down. (Top) Before the onset of P-cell nuclear migration, P cells (grey) span from the lateral to the ventral side of the worm, with the nuclei (pink circles) located laterally. The P cell is partitioned into a lateral and ventral region by a narrow constriction between body wall muscles (orange in cross section) and the cuticle. (Middle) When nuclear migration begins, P-cell nuclei migrate through the constriction towards the ventral region. This process starts with the most anterior pair of P cells and is followed by the consecutive pairs. (Bottom) P-cell nuclear migration ends when all twelve nuclei have migrated to the ventral cord. **(B).** Representative epifluorescence images of L4 animals expressing *unc-47::gfp* in ventral cord GABA neurons in *unc-84(n369)* (top panels) and *unc-84(n369, cgef-1(yc3)* double mutants (bottom panels). Animals expressing an *unc-84(+)* rescue array also express *odr-1::rfp* in the head (left panels). L4 larvae are shown from the lateral side with anterior to the left and ventral down. **(C).** Plot of the number of missing GABA neurons at 15°C, **(D).** 20°C, **(E).** and 25°C of the *unc-84(n369), oxIs12(unc-47::gfp); ycEx60(odr-1::rfp,WRM0617cH07)* and *cgef-1(yc3)* single and double mutants. **(F).** Plot of the number of missing GABA neurons when both *WRM0625dG08* and *WRM0622bA03* fosmids (long fosmids) are expressed at 15°C. Each color represents an independent line with a total of three independent lines assayed for P-cell nuclear migration defects. **(G).** Number of missing GABA neurons when *WRM0627cD01* fosmid (short fosmid) is expressed at 15°C. Each color represents an independent line with a total of four independent lines assayed for P-cell nuclear migration defects. “OE” denotes overexpression of the indicated transgene. All error bars are 95% confidence intervals and all statistical analysis was done using student t-tests. *** indicates a P-value < 0.001 and **** indicates a P-value < 0.0001.

P-cell nuclear migration is regulated by the SUN (**S**ad1 and **UN**C-84) protein UNC-84 and the KASH (**K**larsicht, **A**NC-1, **S**yne **h**omology) protein UNC-83 (Malone et al., 1999; Starr et al., 2001). UNC-84, located at the inner nuclear membrane, interacts with UNC-83 to recruit it to the out nuclear membrane (McGee et al., 2006). Together, UNC-84 and UNC-83 form a LINC (**Li**nker of **N**ucleoskeleton and **C**ytoskeleton) complex to transfer forces generated by the cytoskeleton in the cytoplasm to structures inside the nucleus (Starr & Fridolfsson, 2010). UNC-83 is then able to interact with microtubule motor proteins kinesin-1 and cytoplasmic dynein to move nuclei (Fridolfsson et al., 2010; Fridolfsson & Starr, 2010; Meyerzon et al., 2009). In larval P-cells, dynein is the major motor necessary to move nuclei toward the minus ends of microtubules in the ventral cord (Bone et al., 2016; Ho et al., 2018). Null mutations in *unc-83* or *unc-84* lead to a temperature-sensitive nuclear migration defect in P cells. When these mutants are grown at 25°C, less than 40% of P-cell nuclei successfully migrate to a ventral position. However, when the LINC complex is disrupted at 15°C, at least 90% of P-cell nuclei migrate successfully (Malone et al., 1999; Starr et al., 2001). This leads to our hypothesis that there is an additional pathway that functions parallel to the LINC complex-dependent pathway to move P-cell nuclei through constricted spaces.

To identify players in this alternative nuclear migration pathway, we previously conducted an unbiased forward genetics screen for **e**nhancers of the nuclear **m**igration defect of ***u****nc-84* (*emu*) at 15°C (Chang et al., 2013). Eight *emu* mutations were isolated in these screens and one was identified as a lesion in *toca-1* (Transducer of Cdc-42 dependent actin assembly) (Chang et al., 2013). TOCA-1 is predicted to have an F-BAR domain, a domain that interacts with the Rho GTPase Cdc42, and a domain that interacts with actin-nucleating WASP proteins (Fricke et al., 2009; Giuliani et al., 2009; H.-Y. H. Ho et al., 2004). One model is that TOCA-1 functions by binding to the nuclear membrane, recruiting Cdc42 and WASP to nucleate actin, and deforming the nucleus to aid in migration.

Here we report the identification of a second *emu* allele in *cgef-1* which is predicted to encode a guanine nucleotide exchange factor (GEF) for CDC-42 (Chan & Nance, 2013). GEFs function by activating G-proteins, which are molecular switches involved in regulating signaling cascades. G-proteins can be found in an “inactive” GDP-bound state and an “active” GTP-bound state. GEFs activate G-proteins by facilitating the exchange of GDP for GTP (Rossman et al., 2005; Schmidt & Hall, 2002). The Rho-GTPase family of G-proteins includes RhoA, Rac, and Cdc42, which function by regulating polarity establishment, cell movement, and cytoskeletal dynamics (Etienne-Manneville, 2004; Hall, 1998). *C. elegans* orthologs are RHO-1, CED-10 (Rac), MIG-2 (Rac), and CDC-42 (Reiner & Lundquist, 2018). CGEF-1 acts as GEF for CDC-42 during early embryonic development (Chan & Nance, 2013), but its role outside of embryogenesis is unclear. We propose that CGEF-1 activates CDC-42, which then assembles actin networks to help nuclei migrate through constricted spaces in a pathway that functions parallel to the LINC complex and dynein pathway. To test this hypothesis, we examined the roles of CGEF-1, CDC-42, and other actin regulators during P-cell nuclear migration.

## Results

### Mutations in *cgef-1* enhance the P-cell nuclear migration defect of *unc-84(null)* animals

The *yc3* and *yc21* alleles were found in an *emu* screen and the homozygous mutants significantly enhance the nuclear migration defect of *unc-84* (Chang et al., 2013). To quantify P-cell nuclear migration, we expressed the GABA neuronal marker *p_unc-47_::gfp* and counted the number of GABA neurons at the L4 stage as an indicator of successful P-cell nuclear migration (Fridolfsson et al., 2018; McIntire et al., 1997). At 15°C, *unc-84(n369)* mutants were missing an average of 2.08±0.66 [mean±95% confidence interval (CI)] GABA neurons, slightly above wildtype (Fig. 1B-C). *yc3* single mutants had no phenotype on their own, missing an average of 1.30±0.43 GABA neurons. However, *yc3, unc-84(n369)* double mutants had an average of 4.41±0.73 missing GABA neurons (p<0.00005). *yc3* also enhanced the nuclear migration defect of *unc-84(n369)* at 20°C and at 25°C (Fig. 1D-E).

To identify the molecular lesion underlying *emu* alleles, we performed whole genome sequencing of seven different *emu* mutant strains isolated in our previous screen (Chang et al., 2013). We cataloged single nucleotide polymorphisms (SNPs) that were predicted to cause severe disruptions to open reading frames. SNPs that were found in all the strains were eliminated because they were likely in the background of the UD87 strain that was used for mutagenesis. We focused on a SNP predicted to cause a premature stop codon in the *cgef-1* gene that was identified in both the *yc3* and *yc21* alleles. Nucleotide X:2798063 in the penultimate exon of *cgef-1* was mutated from a G to an A, causing the tryptophan 345 of CGEF-1a to change to a premature stop codon (Fig. 2A).

**Figure 2.**
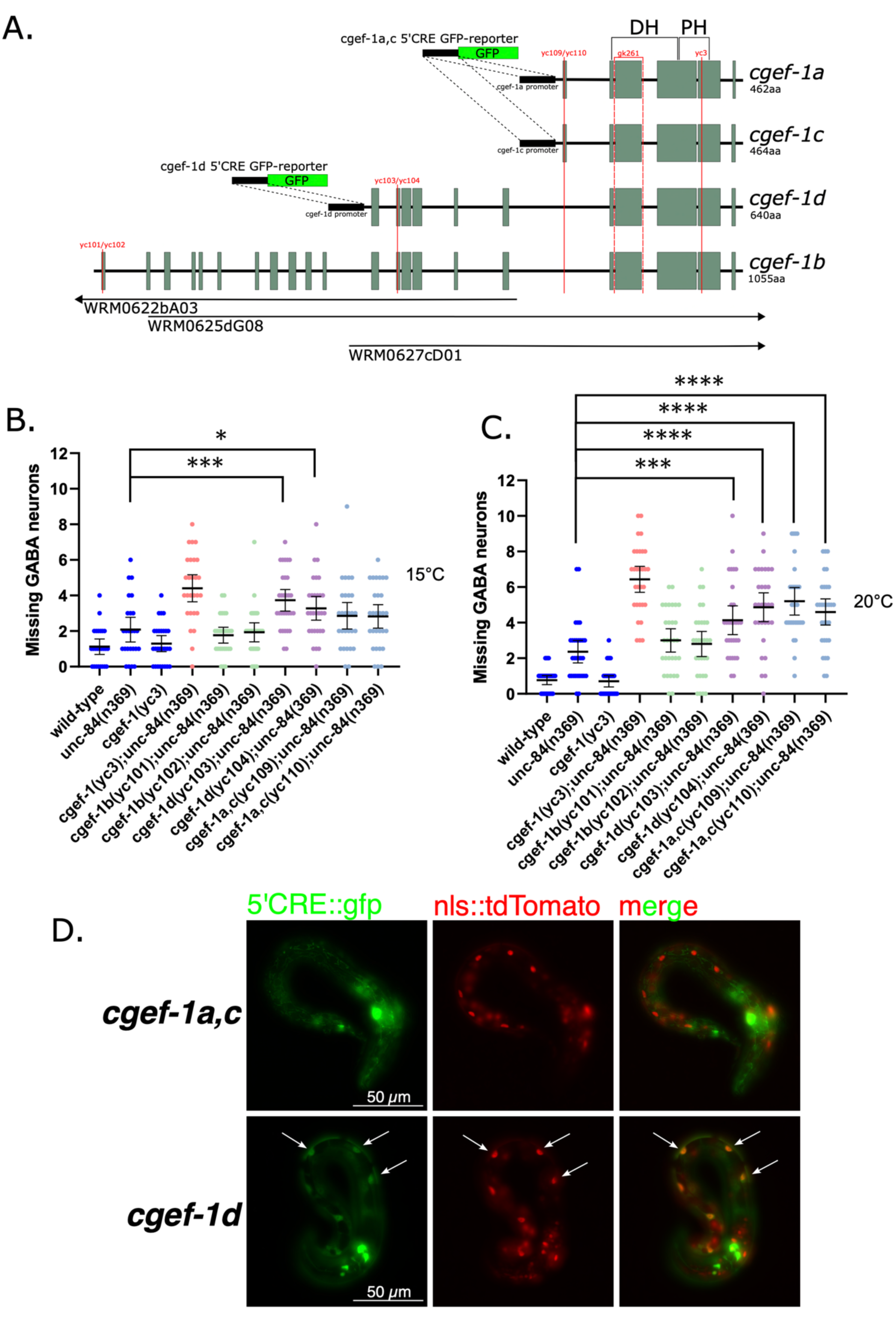
The *cgef-1d* isoform is required in P-cell nuclear migration. **(A).** Intron (black lines) and exon (green boxes) schematics for the four known *cgef-1* isoforms are shown. All isoforms share the same 3’ end, which encodes Dbl homology (DH) and Pleckstrin homology (PH) domains. The red vertical lines that run perpendicular to the diagrams show where the labeled mutations are located. The regions that the *WRM0625dG08, WRM0622bA03,* and *WRM0627cD01* fosmids overlap are indicated at the bottom and the arrows indicate that the fosmids continue beyond the genomic region shown in the diagram. *cgef-1a,c* and *cgef-1d* 5’*cis*-regulatory element (5’CRE) GFP-reporters are shown. These constructs were generated by using 2.5kb region upstream of each isoform transcript to drive GFP expression. **(B).** Plot of the number of missing GABA neurons at 15°C and at **(C).** 20°C of *cgef-1* isoform mutations. **(D).** Representative images of *phlh-3::nls::tdTomato* (red) and *cgef-1a,c* 5’CRE GFP-reporter (top row) and *cgef-1d* 5’CRE GFP-reporter (bottom row). White arrows indicate areas of co-localization. All error bars are 95% confidence intervals and all statistical analysis was done using student t-tests. * indicates a P-value <0.05, *** indicates a P-value < 0.001, and **** indicates a P-value < 0.0001.

To confirm that the premature stop codon in *cgef-1(yc3)* is the molecular lesion responsible for enhancing the nuclear migration defect of *unc-84*, we tested whether other alleles of *cgef-1* also enhance the P-cell nuclear migration defect of *unc-84(n369)* null mutants. *cgef-1(gk261)* is likely a null allele, as it is a 318 bp deletion that removes the entire third exon in *cgef-1a* and is predicted to result in a frame shift (Fig. 2A) (*C. elegans* Deletion Mutant Consortium, 2012). *cgef-1(gk261) unc-84(n369)* double mutants had significant P-cell nuclear migration defects at 15°C, 20°C, and 25°C compared to the single mutants (Fig 1C-E). Likewise, *cgef-1(RNAi)* significantly enhanced the nuclear migration defects of *unc-84(n369)* (Fig. 1C-E). Thus, multiple alleles and RNAi of *cgef-1* all had similar phenotypes.

To confirm that the *yc3* phenotype is due to the molecular lesion in *cgef-1* and not some other mutation in the genome, we expressed *cgef-1* extra-chromosomal rescue arrays in *yc3, unc-84(n369)* double mutants. A fosmid covering the longest isoforms of *cgef-1*(WRM0622bA03) was able to rescue the nuclear migration defect in three independent lines (P<0.0005) (Fig. 1F). Together, these data strongly suggest that lesions in *cgef-1* are responsible for the nuclear migration defects observed in *yc3* and *yc21* animals and that *cgef-1* functions in parallel with the LINC complex to facilitate P-cell nuclear migration.

### The *cgef-1d* isoform is necessary for LINC complex-dependent P-cell nuclear migration

CGEF-1 activates CDC-42 during early embryogenesis (Chan & Nance, 2013; Kumfer et al., 2010). *C. elegans* encodes at least four *cgef-1* predicted isoforms (Fig. 2A) (Ziel et al., 2009; Wormbase). All four isoforms share the last four exons of the *cgef-1* gene, which are predicted to encode a catalytic Dbl homology (DH) domain as well as a pleckstrin homology (PH) domain (Chan & Nance, 2013; Ziel et al., 2009). *cgef-1a* and *cgef-1c* encode the shortest CGEF-1 isoforms that differ in length by only two amino acids at their N termini. *cgef-1b* encodes the longest CGEF-1 isoform, while *cgef-1d* encodes an isoform that is intermediate in length. Expression of a fosmid (WRM0627cD01) that only spans the *cgef-1a, c,* and *d* isoforms was sufficient to rescue the *cgef-1(yc3) unc-84* nuclear migration defect at 15°C, indicating that exons 1 – 11 of *cgef-1b* are not necessary for P-cell nuclear migration (Fig. 1G).

To further determine which isoform of *cgef-1* is necessary for this process, we generated new alleles in each isoform using CRISPR-Cas9 gene editing, either as an early stop codon in the first or second exon of an isoform or as a deletion mutation that resulted in a predicted frameshift (Fig. 2A). Predicted severe alleles of the long isoforms, *cgef-1b(yc101)* and *cgef-1b(yc102)*, did not enhance the nuclear migration defect of *unc-84(n369)* at 15 or 20°C (Fig. 2B-C). In contrast, the *cgef-1d(yc103)* early stop codon and *cgef-1d(yc104)* frame-shift deletions mutations significantly enhanced the nuclear migration defect of *unc-84(n369)* at 15 and 20°C (Fig. 2C). Mutations in the shortest isoforms, *cgef-1a,c(yc109)* and *cgef-1a,c(yc110)*, also enhanced the nuclear migration defect of *unc-84(n369)* at 20°C but not at 15°C. Thus, we conclude that *cgef-1b* is dispensable for P-cell nuclear migration while *cgef-1d* and *cgef-1a,c* are necessary for nuclear migration when the LINC complex is disrupted.

We next determined whether the *cgef-1a,c* and *cgef-1d* isoforms are expressed in P-cells during nuclear migration. We used previously described 5’*cis*-regulatory element reporter strains that drive the expression of GFP with a nuclear localization signal under control of promoters for or *cgef-1d* or *cgef-1a,c* (Ziel et al., 2009) (Fig. 2A) and looked for GFP expression in L1 larval P-cell nuclei, which were marked with a tdTomato nuclear marker expressed from the P-cell specific promoter of *hlh-3* (Bone et al., 2016; Chang et al., 2013). The *cgef-1d* reporter expressed GFP in larval P cells. However, the *cgef-1a,c* reporter did not express detectable GFP above background in larval P cells. Thus, *cgef-1d* appears to be the main isoform expressed in P-cells and it is necessary for LINC complex-dependent nuclear migration in these cells.

### CGEF-1 activates CDC-42 during P-cell nuclear migration

Since CGEF-1 activates the small GTPase CDC-42 in early embryogenesis (Chan & Nance, 2013; Kumfer et al., 2010), we hypothesized that *cdc-42* is downstream of *cgef-1* and is necessary for P-cell nuclear migration in the absence of *unc-84*. To test this, we knocked-down *cdc-42* specifically in larval P cells at the time of nuclear migration using the auxin-inducible degradation (AID) system (J. Ho et al., 2018; Zhang et al., 2015). We tagged the endogenous *cdc-42* locus with a 44-amino acid degron using CRISPR/Cas9 engineering and expressed the TIR-1 E3 ubiquitin ligase under control of the P-cell specific *hlh-3* promoter. This combination of tissue-specific expression of TIR-1 and the addition of auxin during the mid-L1 larval stage, when P-cell nuclear migration occurs, allowed for spatial and temporal control of CDC-42 protein degradation. We found that degrading CDC-42 in otherwise wild-type L1 larvae had no effect on P-cell nuclear migration (Fig. 3A). However, CDC-42 auxin-induced degradation significantly enhanced the *unc-84(null)* nuclear migration defect at both 15 and 25°C (Fig. 3A-B), suggesting that CDC-42 is necessary for P-cell nuclear migration in the absence of LINC complexes.

**Figure 3:**
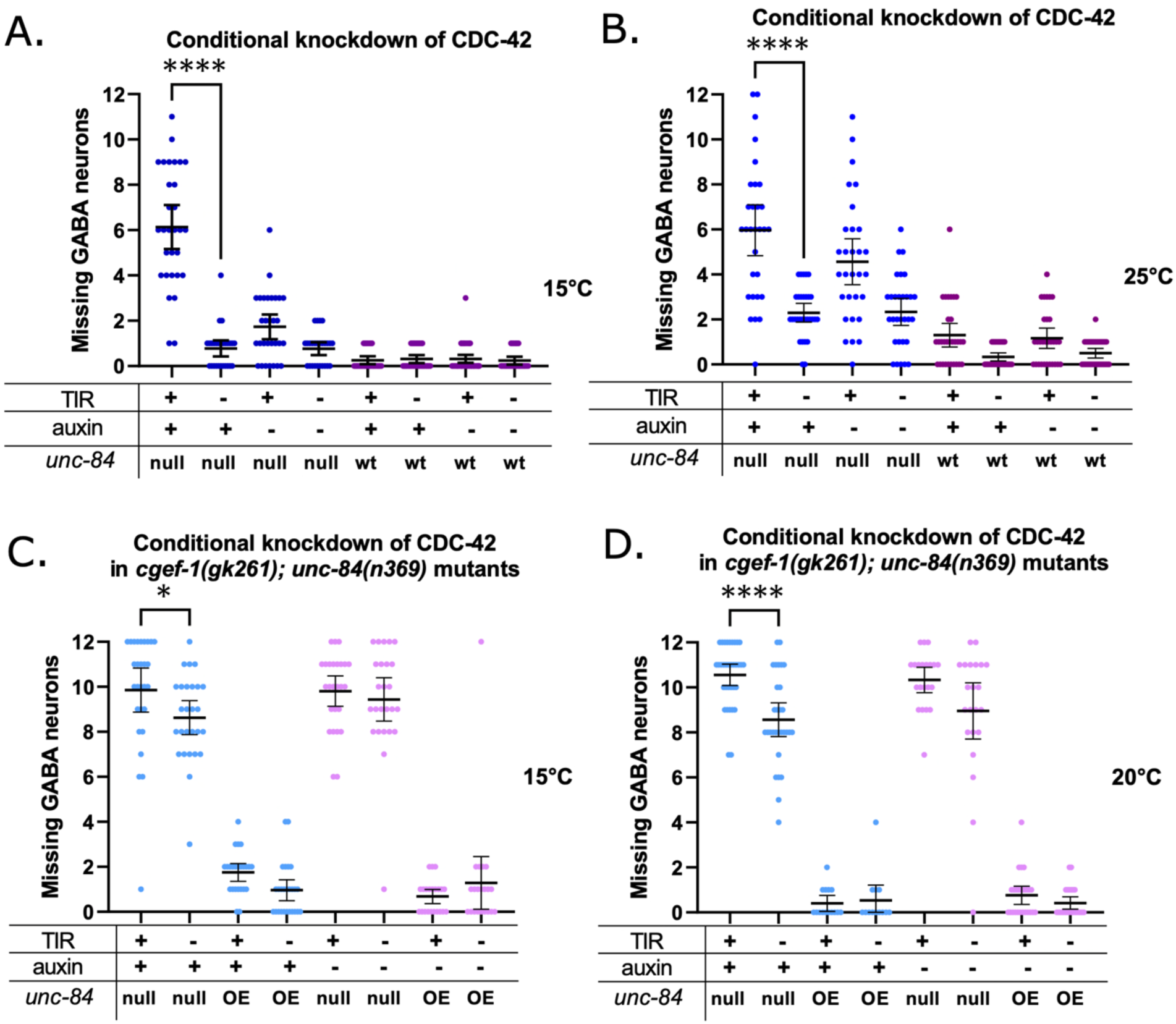
CDC-42 is necessary for P-cell nuclear migration. **(A).** Auxin inducible degradation (AID) system was used to knock down CDC-42 in P cells at 15C° and **(B).** 25C°. *degron::GFP11::cdc-42; unc-84(n369), oxIs12(unc-47::gfp); ycEx266(Pmyo-2::mCherry;phlh-3::TIR-1::mRuby)* shown in blue dots and *degron::GFP11::cdc-42; oxIs12(unc-47::gfp); ycEx266(Pmyo-2::mCherry;phlh-3::TIR-1::mRuby)* shown in purple dots were treated with auxin during P-cell nuclear migration. The y-axis shows the number of missing GABA neurons. **(C).** AID was used to knockdown CDC-42 in P cells of *degron::GFP11::cdc-42*; *unc-84(n369), cgef-1(gk261), oxIs12(unc-47::gfp); ycEx300(odr-1::gfp,WRM0617cH07); ycEx266(Pmyo-2::mCherry;phlh-3::TIR-1::mRuby)* mutants at 15C° and at **(D).** 20C°. Light blue dots indicate strains that were exposed to auxin and pink dots indicate strains that were not exposed to auxin. For all the graphs, “OE” and “null“ for *unc-84* indicates *unc-84(n369)* null mutation with expression of the *unc-84* rescue array (*ycEx60(odr-1::rfp,WRM0617cH07))* and no expression of the rescue array, respectively. “wt” indicates no *unc-84(n369)* mutation. All error bars are 95% confidence intervals and all statistical analysis was done using student t-tests. **** indicates a P-value < 0.0001.

We next tested whether *cdc-42* is in the same pathway as *cgef-1* by degrading CDC-42 in *cgef-1, unc-84* double mutants. Degradation of CDC-42 in *cgef-1(gk261), unc-84(n369)* double mutant L1 larvae significantly enhanced the P-cell nuclear migration defects of the double mutant alone (Fig. 3C-D). Because *cgef-1(gk261)* is a predicted null, this result suggests that CGEF-1 and CDC-42 have partially independent roles and that other RhoGTPases or GEFs may function during P-cell nuclear migration.

To test the roles of other small RhoGTPases that might function downstream of *cgef-1*, we expressed constitutively active *cdc-42, rho-1, mig-2,* or *ced-10* (Alan et al., 2013; Gujar et al., 2019; Norris et al., 2014) to see if they suppressed the nuclear migration defects of *cgef-1(yc3), unc-84(n369)* double mutants. When constitutively active *cdc-42(G12V)* was expressed from an extrachromosomal array under control of the P-cell specific *hlh-3* promoter, it was able to partially rescue the nuclear migration defect of *cgef-1(yc3), unc-84(n369)* double mutants (Fig. 4A-B). Thus, we conclude that *cdc-42* is downstream of and likely activated by *cgef-*1. However, when constitutively active *rho-1(G14V), mig-2(G16V),* or *ced-10(G12V)* were expressed in P cells, we did not observe rescue of nuclear migration defects in *cgef-1(yc3), unc-84(n369)* mutants. Instead, we observed an enhancement of P-cell nuclear migration defects when Rac family members *mig-1(G16V)* or *ced-10(G12V)* were overexpressed in *cgef-1(yc3), unc-84(n369)* mutants (Fig. 4C-E), consistent with previous findings that Rac functions during P-cell migration (Spencer et al 2001). In summary, we conclude that CGEF-1 activates CDC-42 during P-cell nuclear migration.

**Figure 4:**
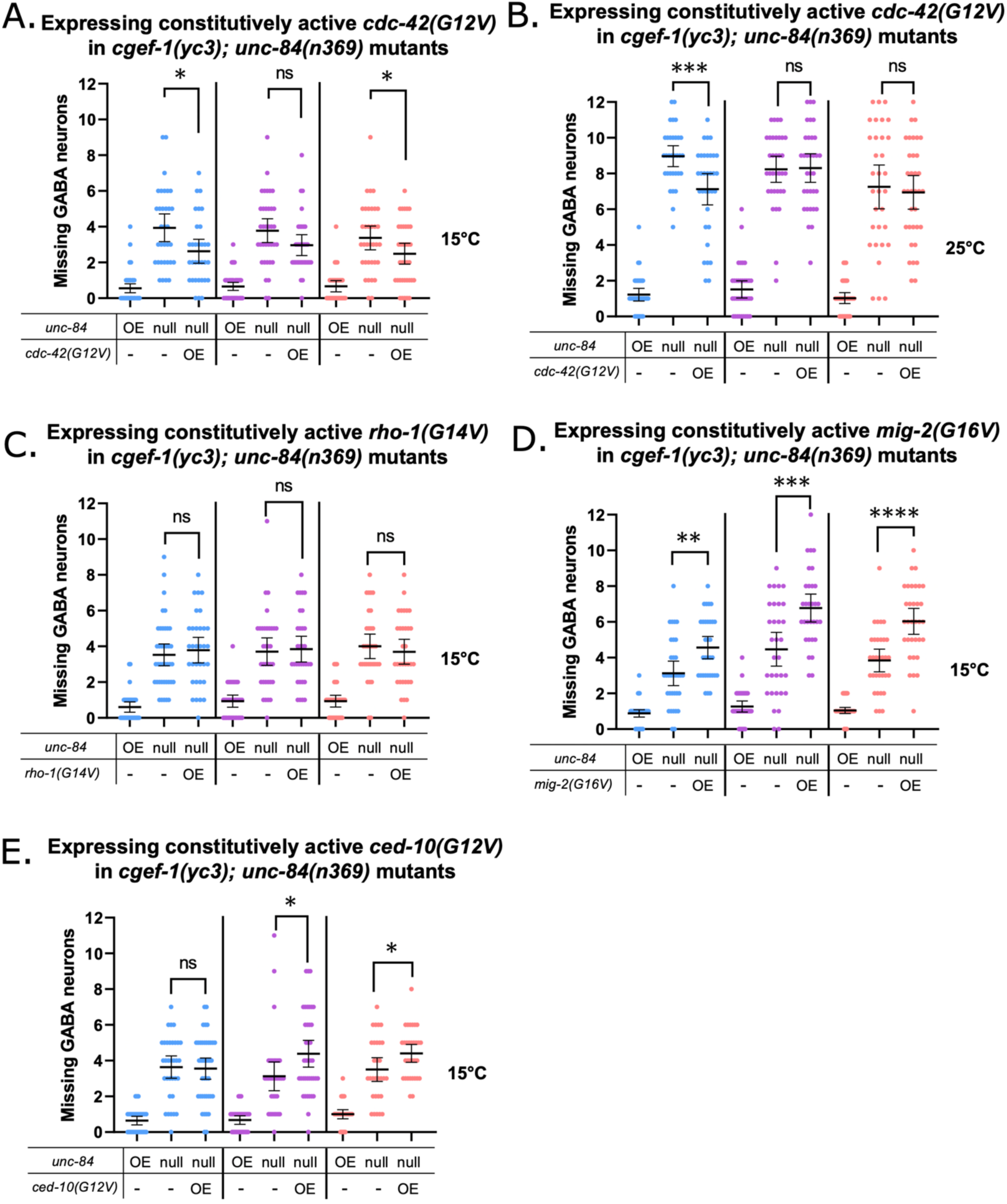
CGEF-1 activates CDC-42 during P-cell nuclear migration. **(A).** *phlh-3::2xHA::cdc-42(G12V); odr-1::gfp* construct was expressed in *unc-84(n369);cgef-1(yc3);oxIs12(unc-47::gfp);ycEx60(odr-1::rfp,WRM0617cH07)* in three independent lines as indicated by the different colors. Worms were grown at 15C° and **(B).** 25C° and assayed for P-cell nuclear migration. The y-axis shows the number of missing GABA neurons. **(C).** *phlh-3::2xHA::rho-1(G14V); odr-1::gfp*, **(D).** *phlh-3::2xHA::mig-2(G16V); odr-1::gfp*, **(E).** and *phlh-3::2xHA::ced-10(G12V); odr-1::gfp* constructs were expressed in *unc-84(n369);cgef-1(yc3);oxIs12(unc-47::gfp);ycEx60(odr-1::rfp,WRM0617cH07)* in three independent lines as indicated by the different colors. Worms were grown at 15C° and assayed for P-cell nuclear migration. The y-axis shows the number of missing GABA neurons. For all the graphs, “OE” and “null“ for *unc-84* indicates expression of the *unc-84* rescue array (*ycEx60(odr-1::rfp,WRM0617cH07))* and no expression of the rescue array, respectively. “OE” for the other transgene indicate overexpression of the indicated transgene. All error bars are 95% confidence intervals and all statistical analysis was done using student t-tests. * indicates a P-value <0.05, ** indicates a P-value < 0.01, *** indicates a P-value < 0.001, and **** indicates a P-value < 0.0001.

### Branched actin and actin-myosin networks are necessary for P-cell nuclear migration

While CDC-42 is necessary for nuclear migration, it is not clear how it functions. CDC-42 regulates many downstream effectors. Here we tested potential CDC-42 effectors involved in cell polarity (PAR-6 and PKC-3), actin networks (ARX-3 of the Arp2/3 complex), and actomyosin contractions (NMY-2), to determine their necessity for P-cell nuclear migration.

CDC-42 activates PAR-6 and PKC-3 during polarization events during early embryogenesis (Aceto et al., 2006; Gotta et al., 2001; Wang et al., 2017) and regulates non-centrosomal microtubule arrays in the larval epidermal epithelium (Castiglioni et al., 2020). We therefore hypothesized that CDC-42 functions through PAR-6 and PKC-3 during P-cell nuclear migration. We used the AID system to knock down PAR-6 and PKC-3 in P cells during nuclear migration (Castiglioni et al., 2020). Degradation of PAR-6 slightly enhanced the *unc-84(n369)* nuclear migration defect at 15C° but not at 20C° (Fig. 5A-B). However, this defect was mild compared to degrading CDC-42 in *unc-84(n369)* mutants. Furthermore, the degradation of PKC-3 did not significantly enhance the nuclear migration defect of *unc-84(n369)* (Fig. 5C-D). Together, these data suggest that PAR-6 plays only a minor role during P-cell nuclear migration and that and CDC-42 is functioning through an alternative pathway.

**Figure 5:**
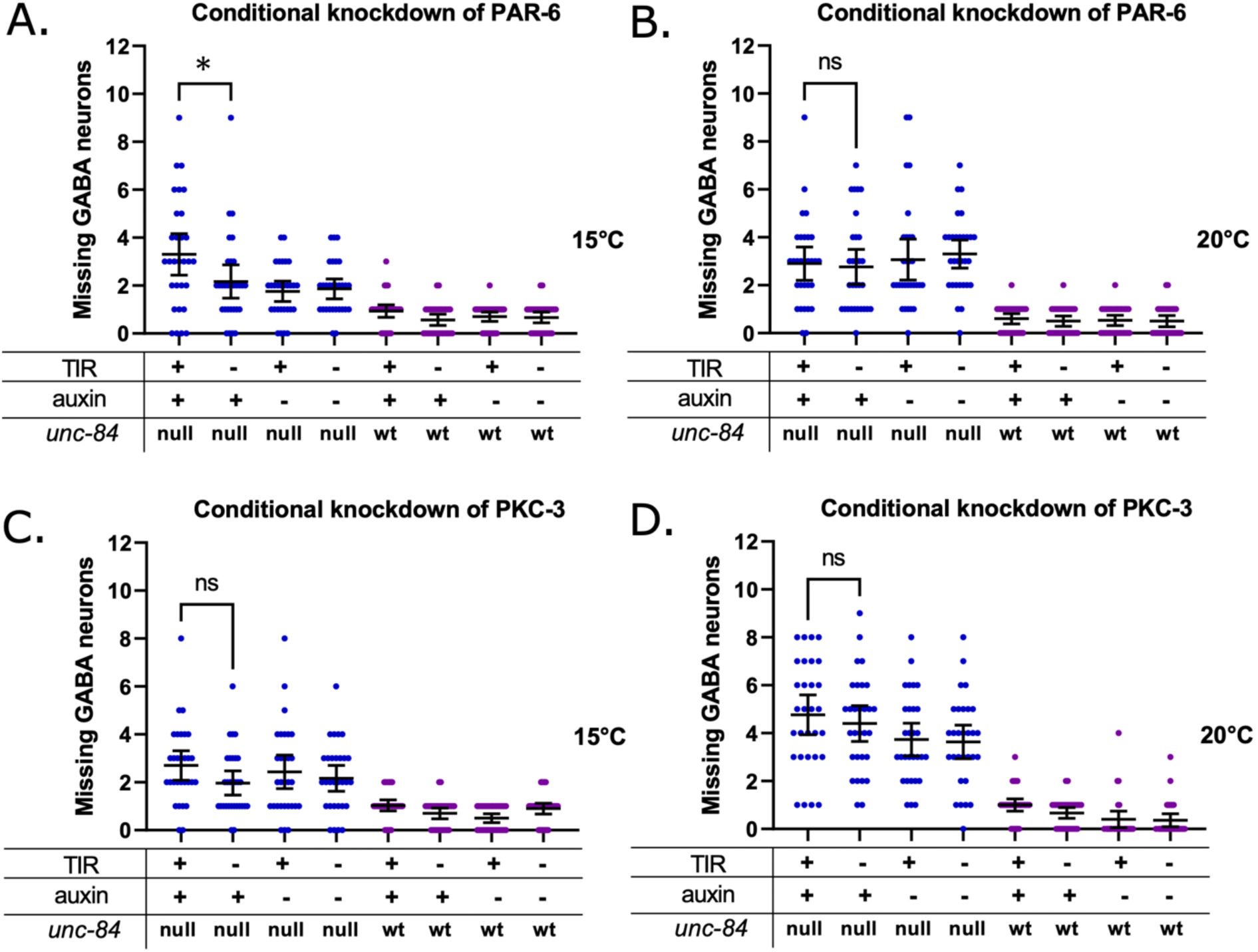
PAR-6 and PKC-3 are not involved in P-cell nuclear migration. **(A).** AID was used to knock down PAR-6 in P cells at 15C° and **(B).** 20C°. *par-6::degron::egfp; unc-84(n369), oxIs12(unc-47::gfp); ycEx266(Pmyo-2::mCherry;phlh-3::TIR-1::mRuby)* shown in blue dots and *par-6::degron::egfp; oxIs12(unc-47::gfp); ycEx266(Pmyo-2::mCherry;phlh-3::TIR-1::mRuby)* shown in purple dots were treated with auxin during P-cell nuclear migration. The y-axis shows the number of missing GABA neurons. **(C).** AID was used to knock down PKC-3 in P cells at 15C° and **(D).** 20C°. *egfp::degron::pkc-3; unc-84(n369), oxIs12(unc-47::gfp); ycEx266(Pmyo-2::mCherry;phlh-3::TIR-1::mRuby)* shown in blue dots and *egfp::degron::pkc-3; oxIs12(unc-47::gfp); ycEx266(Pmyo-2::mCherry;phlh-3::TIR-1::mRuby)* shown in purple dots were treated with auxin during P-cell nuclear migration. The y-axis shows the number of missing GABA neurons. For all the graphs, “OE” and “null“ for *unc-84* indicates *unc-84(n369)* null mutation with expression of the *unc-84* rescue array (*ycEx60(odr-1::rfp,WRM0617cH07))* and no expression of the rescue array, respectively. “wt” indicates no *unc-84(n369)* mutation. All error bars are 95% confidence intervals and all statistical analysis was done using student t-tests. * indicates a P-value <0.05

A major function performed by CDC-42 is the regulation of the assembly and dynamics of actin networks (Carlier et al., 1999; Ma et al., 1998; Rohatgi et al., 1999). To test the hypothesis that branched-actin networks function during P-cell nuclear migration, we used the AID system to degrade a component of the Arp2/3 complex. ARX-3, the *C. elegans* homolog of mammalian Arp3, is one of the seven subunits that make up the ARP2/3 complex (Sawa et al., 2003). Degradation of ARX-3 in L1 larvae at the time of P-cell nuclear migration had no defect on its own, again supporting the hypothesis that the LINC complex pathway is sufficient to move P-cell nuclei (Fig. 6A-B). However, degrading ARX-3 significantly enhanced the *unc-84(n369)* nuclear migration defect (Fig. 6A-B). Thus, we conclude that the Arp2/3 complex is necessary for the movement of P-cell nuclei in the absence of LINC complexes.

**Figure 6:**
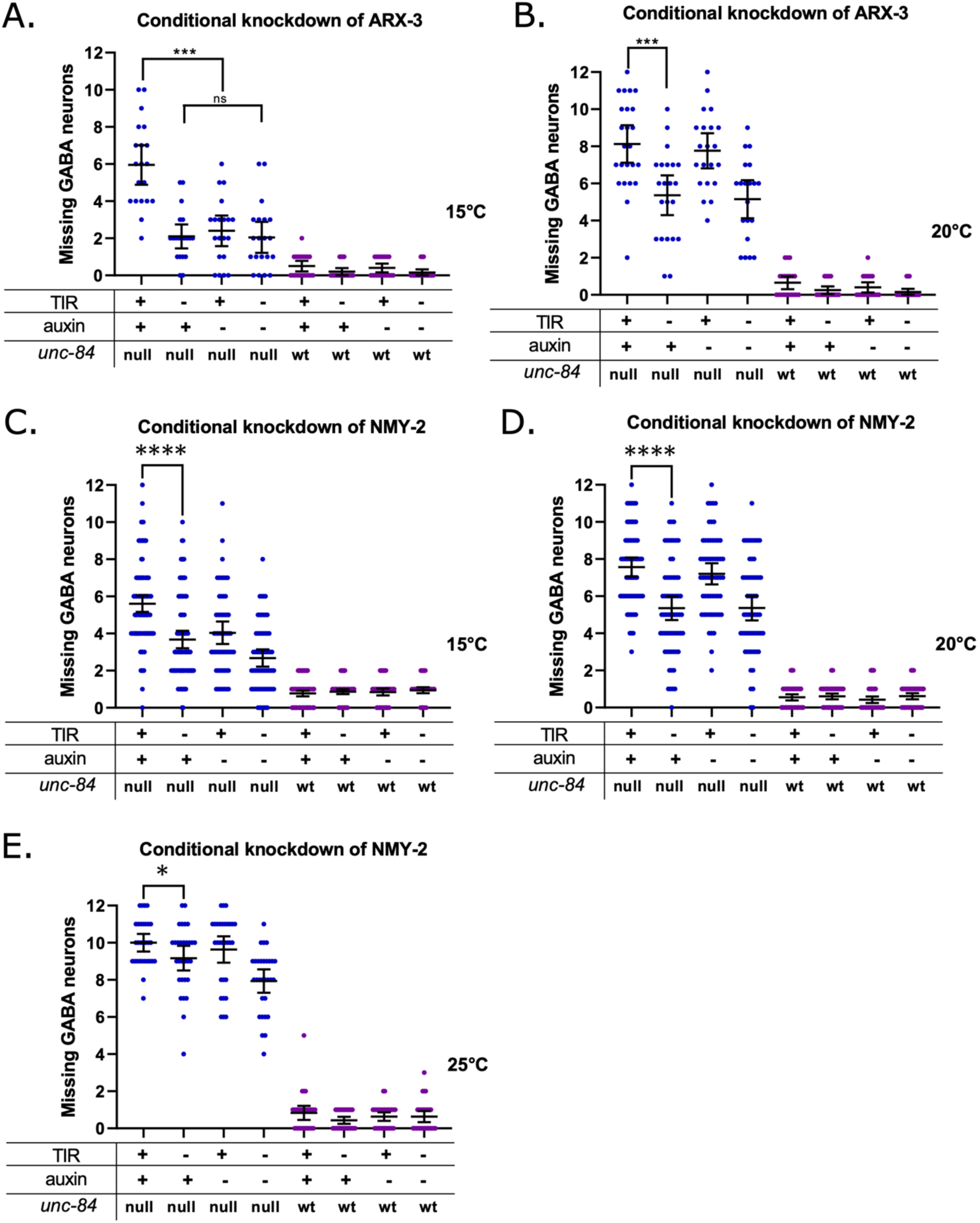
Branched actin and actin-myosin contractions are involved in P-cell nuclear migration. **(A-B).** Auxin inducible degradation system was used to knock down ARX-3 in P cells at 15°C (A) or 20°C (B). *degron::arx-3; unc-84(n369), oxIs12(unc-47::gfp); ycEx253(odr-1::rfp;phlh-3::TIR-1::mRuby)* shown in blue dots and *degron::arx-3; oxIs12(unc-47::gfp); ycEx253(odr-1::rfp;phlh-3::TIR-1::mRuby)* shown in purple dots were treated with auxin during P-cell nuclear migration. The y-axis shows the number of missing GABA neurons. **(C).** Auxin inducible degradation system was used to knock down NMY-2 in P cells at 15C°, **(D).** 20C°, and **(E).** 25C°. *degron::GFP11::nmy-2; unc-84(n369), oxIs12(unc-47::gfp); ycEx266(Pmyo-2::mCherry;phlh-3::TIR-1::mRuby)* shown in blue dots and *degron::GFP11::nmy-2; oxIs12(unc-47::gfp);ycEx266(Pmyo-2::mCherry;phlh-3::TIR-1::mRuby)* shown in purple dots were treated with auxin during P-cell nuclear migration. The y-axis shows the number of missing GABA neurons. For all the graphs, “OE” and “null“ for *unc-84* indicates *unc-84(n369)* null mutation with expression of the *unc-84* rescue array (*ycEx60(odr-1::rfp,WRM0617cH07))* and no expression of the rescue array, respectively. “wt” indicates no *unc-84(n369)* mutation. All error bars are 95% confidence intervals and all statistical analysis was done using student t-tests. * indicates a P-value <0.05, *** indicates a P-value < 0.001, and **** indicates a P-value < 0.0001.

We next hypothesized that myosin may be working with actin networks to exert pushing or pulling forces on P-cell nuclei. To test this hypothesis, we used the AID system by adding a degron tag onto the N terminus of the non-muscle myosin heavy chain *(nmy-2)* gene. Degradation of NMY-2 in an *unc-84(n369)* background resulted in a significantly worse nuclear-migration defect than the *unc-84(n369)* single mutant larvae (Fig. 6C-E). Therefore, CDC-42, ARX-3, and NMY-2 are all necessary to migrate P-cell nuclei in the absence of LINC complexes.

## Discussion

Our findings presented here support that two parallel pathways facilitate P-cell nuclear migration: a LINC complex-dependent pathway and an actin-dependent pathway (Fig. 7). For the LINC complex pathway, the SUN protein UNC-84 spans the inner nuclear membrane and interacts with the KASH protein UNC-83 in the outer nuclear membrane. The cytoplasmic domain of UNC-83 then interacts with microtubule motors kinesin-1 and dynein (Fridolfsson et al., 2010; Fridolfsson & Starr, 2010; Meyerzon et al., 2009). In P-cell nuclear migration, UNC-83 specifically recruits dynein to the nuclear envelope where it is the main motor protein to pull nuclei towards the minus ends of microtubules (Bone et al., 2016; Ho et al., 2018).

**Figure 7:**
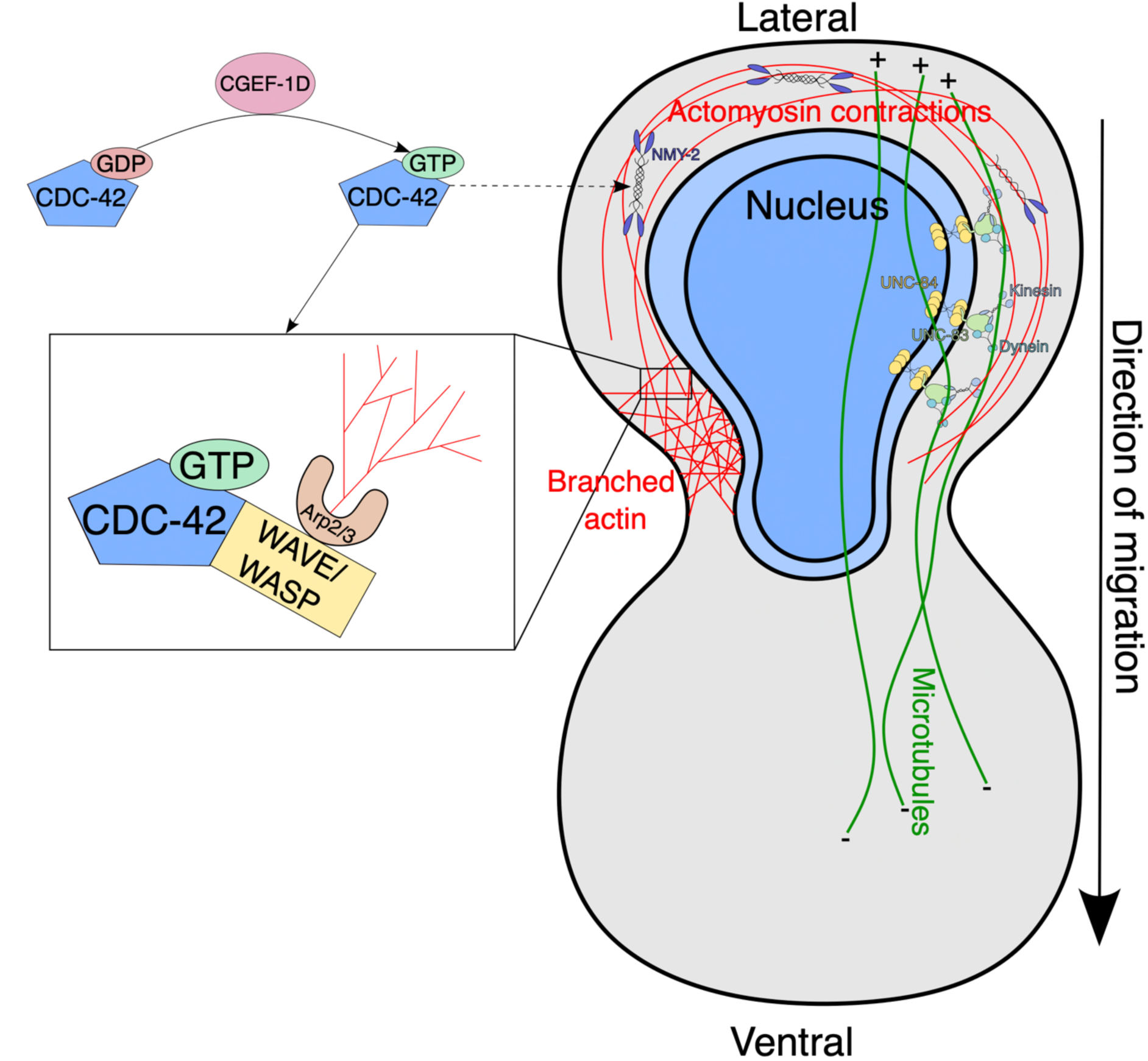
Model for both pathways involved in P-cell nuclear migration. In the LINC complex-dependent pathway (right side of cell), UNC-84 (yellow) interacts with UNC-83 (light green) at the nuclear envelope. UNC-83 interacts with both kinesin and dynein but uses dynein as the main microtubule motor protein to pull the nucleus towards the minus ends of microtubules (green lines). The actin-based pathway is drawn on the left of the cell. In the inset, CGEF-1D activates CDC-42 by exchanging the nucleotide for GTP. CDC-42 in the GTP form activates WAVE/WASP, which then activates the Arp2/3 complex to generate branched actin. CDC-42 may also be involved in indirectly activating NMY-2 which can then induce actomyosin contractions. See text for more details of our proposed model.

Here, we found that an actin-dependent pathway is necessary for P-cell nuclear migration in the absence of the LINC complex pathway. In our model, CGEF-1D is a GEF, which activates the small GTPase CDC-42 during P-cell nuclear migration. CDC-42 then activates the Arp2/3 complex to nucleate branched actin. TOCA-1 and WAVE/WASP likely also work at the level of CDC-42 and Arp2/3 in this pathway (Chang et al., 2013; Giuliani et al., 2009; Raduwan et al., 2020). Once Arp2/3 nucleates actin, other proteins organize actin into networks. We found that NMY-2 is also necessary in the absence of the LINC complex-dependent pathway. One hypothesis is that NMY-2 provides forces to contract actin networks during nuclear migration. Although it is known that CDC-42 can indirectly regulate NMY-2 (Gally et al., 2009; Kumfer et al., 2010; Raduwan et al., 2020; Ramesh et al., 1997; Rohatgi et al., 1999, 2000; Watson et al., 2017), further studies will need to be done to determine if and how CDC-42 regulates NMY-2 in this context.

While our genetic findings implicate actin networks in nuclear migration, it is still not clear where in the cell actin and/or actomyosin specific structures are required to move nuclei. One possibility is that branched actin could be localized at the leading edge of the nucleus to deform it as it enters the constriction. This would be analogous to nuclei in mouse dendritic cells induced to migrate through fabricated constrictions (Thiam et al., 2016). Alternatively, actomyosin contractions could localize to the back of P-cell nuclei to provide a pushing force in the direction of migration. In migrating mammalian neurons, actomyosin contraction behind the nucleus pushes it forward and into constricted spaces (Tsai et al., 2007). In support of this model, there are thick actin cables along the direction of migration and some cells have actin rings on the lateral side of the cell near the trailing end of the nucleus during P-cell nuclear migration (Bone et al., 2016). Actomyosin contractions also provide the force that is needed for dendritic cells to migrate through confined environments with myosin enrichment at the cell rear during contraction (Barbier et al., 2019; Lämmermann et al., 2008). Future studies are required to better understand where in the cell actin-dependent pathways function during nuclear movements.

One advantage of *C. elegans* P cells as a model is that the formation of cellular protrusions, often associated with cell migrations, can be genetically separated from nuclear migrations through constricted spaces. During embryonic development, hypodermal P cells extend cytoplasmic protrusions from the lateral side of the embryo; these protrusions meet at the ventral midline to cover up the endoderm during embryonic ventral enclosure (Williams-Masson et al., 1997). Later in larval development, P-cell nuclei migrate through constrictions from lateral to ventral positions as described above (Sulston & Horvitz, 1977). Different Rho-family GTPases function at different stages of P-cell development. Unlike in LINC complex or *cdc-42* mutants described in this paper, P-cell nuclei in *rho-1* and *ced-10; mig-2* (Rac) mutants remain at their lateral starting points, retract their cytoplasm into the lateral region, remain alive, and form ectopic GABA neurons and pseudovulvae in the lateral side of the animal (Spencer et al., 2001). Finally, even in the worst phenotypes reported here, when both the LINC complex-dependent and actin-dependent pathways were knocked out, some P-cell nuclei still successfully migrated, suggesting there are other pathways yet to be elucidated. Thus, P-cells will continue to be a valuable model for studying nuclear migrations through constricted spaces in development.

## Materials and Methods

### Whole-genome sequencing of strains from the enhancer of the nuclear migration defect of unc-84 screen

Strains carrying the *yc3*, *yc15*, *yc16*, *yc18*, *yc20*, and *yc21* alleles isolated from a previously described chemical mutagenesis screen for *enhancers of the nuclear migration defect of unc-84 (emu)* (Chang et al., 2013). We collected genomic DNA from each homozygous mutant strain for whole-genome sequencing to identify candidate lesions underlying the nuclear migration phenotypes. Genomic DNA preps were made with the Qiagen DNeasy Blood and Tissue kit as previously described (Herrera & Starr, 2018). Genomic DNA was fragmented and made into libraries for Illumina HiSeq2500 sequencing by the Functional Genomics Laboratory at UC Berkeley. RAPID Sequencing where 150 bp PE reads were generated. We processed raw reads using the default settings of the CloudMap pipeline for Galaxy (Minevich et al., 2012). For each mutant line, we generated a list of variants that did not match the reference N2 genome. We excluded variants found in common between mutant lines as they were likely to be variants present in our starting strain used for the mutagenic screen (Doitsidou et al., 2016). We focused on early stop codon mutations that were present in just one or two of the six sequenced strains and identified a single nucleotide polymorphism (SNP) in the *yc3* and *yc21* strains but not the other four sequenced strains. The SNP was predicted to cause an early stop codon in *cgef-1*. No other unique mutations predicted to cause stop codons were identified in the six strains.

### *C. elegans* strains and genetics

*C. elegans* animals were grown on NGM plates seeded with OP50 at their specified temperatures (Brenner, 1974). Some *C. elegans* strains used in this study were provided by the Caenorhabditis Genetics Center (CGC) which is funded by the National Institutes of Health Office of Research Infrastructure Programs (P40OD010440). The strains used in this study are described in Table 1.

**Table 1.** (Strains)

| Strain | Genotype | Reference |
| --- | --- | --- |
| Figure 1 |  |  |
| N2 | Bristol, wild type | (Brenner, 1974) |
| EG1285 | *oxIs12[p<sub>unc-47</sub>::gfp] X* | (McIntire et al., 1997) |
| UD87 | *unc-84(n369), oxIs12 X; ycEx60[odr-1::rfp, WRM0617cH07]* | (Chang et al., 2013) |
| UD279 | *cgef-1(yc21), unc-84(n369), oxIs12 X; ycEx60* | (Chang et al., 2013) |
| UD285 | *cgef-1(yc3), unc-84(n369), oxIs12 X; ycEx60* | (Chang et al., 2013) |
| VC506 | *cgef-1(gk261) X* | (Kumfer et al., 2010) |
| UD923 | *cgef-1(gk261), unc-84(n369), oxIs12 X; ycEx60* | This study |
| UD524 | *cgef-1(yc3), unc-84(n369), oxIs12 X; ycEx60; ycEx250[100 ng/ul odr-1::gfp; 5 ng/ul WRM0625dG08; 5 ng/ul WRM0622bA03]* | This study |
| UD525 | *cgef-1(yc3), unc-84(n369), oxIs12 X; ycEx60; ycEx250[100 ng/ul odr-1::gfp; 5 ng/ul WRM0625dG08; 5 ng/ul WRM0622bA03]* | This study |
| UD526 | *cgef-1(yc3), unc-84(n369), oxIs12 X; ycEx60; ycEx250[100 ng/ul odr-1::gfp; 5 ng/ul WRM0625dG08; 5 ng/ul WRM0622bA03]* | This study |
| UD718 | *cgef-1(yc3), unc-84(n369), oxIs12 X; ycEx60; ycEx267[100ng/uL odr-1; 5ng/uL WRM0627cD01]* | This study |
| UD719 | *cgef-1(yc3), unc-84(n369), osIs12 X; ycEx60; ycEx268[100ng/uL odr-1; 5ng/uL WRM0627cD01]* | This study |
| UD720 | *cgef-1(yc3), unc-84(n369), osIs12 X; ycEx60; ycEx269[100ng/uL odr-1; 5ng/uL WRM0627cD01]* | This study |
| UD721 | *cgef-1(yc3), unc-84(n369), oxIs12 X; ycEx60; ycEx270[100ng/uL odr-1; 5ng/uL WRM0627cD01]* | This study |
| Figure 2 |  |  |
| UD814 | *cgef-1b(yc101[S14\*]), unc-84(n369), oxIs12 X; ycEx60* | This study |
| UD815 | *cgef-1b(yc102[Δ50 nt in exon 1]), unc-84(n369), oxIs12 X; ycEx60* | This study |
| UD818 | *cgef-1d(yc103[G13\*]), unc-84(n369), oxIs12 X; ycEx60* | This study |
| UD819 | *cgef-1d(yc104[Δ49 nt in exon 2]), unc-84(n369), oxIs12 X; ycEx60* | This study |
| UD822 | *cgef-1a,c(yc109[P6\*]), unc-84(n369), oxIs12 X; ycEx60* | This study |
| UD823 | *cgef-1a,c(yc110[Δ52 nt exon 1]), unc-84(n369), oxIs12 X; ycEx60* | This study |
| NK774 | *qyEx116 [p<sub>cgef-1b</sub>::GFP + unc-119(+)]* | (Ziel et al., 2009) |
| NK775 | *qyEx117 [p<sub>cgef-1a,c</sub>::GFP + unc-119(+)]*. | (Ziel et al., 2009) |
| UD725 | *qyEx116[p<sub>cgef-1b</sub>::GFP + unc-119(+)]*; *ycEx244 [odr-1::gfp; p<sub>hlh-3</sub>::nls::tdTomato]* | This study |
| UD726 | *qyEx117[p<sub>cgef-1a,c</sub>::GFP + unc-119(+)]*; *ycEx244 [odr-1::gfp; p<sub>hlh-3</sub>::nls::tdTomato]* | This study |
| Figure 3 |  |  |
| AFS222 | *zen-4(cle10) IV* | (Farboud et al., 2019) |
| UD716 | *cdc-42(yc100[degron::GFP11::cdc-42]) II*; *oxIs12 X*; *ycEx266[p<sub>myo-2</sub>::mCherry; p<sub>hlh-3</sub>::TIR-1::mRuby]* | This study |
| UD717 | *cdc-42(yc100) II*; *unc-84(n369)*, *oxIs12 X*; *ycEx266* | This study |
| UD989 | *cdc-42(yc100[degron::GFP11::cdc-42]) II*; *cgef-1(gk261)*, *unc-84(n369)*, *oxIs12 X*; *ycEx300[odr-1::GFP, unc-84 (+)]*; *ycEx266[p<sub>myo-2</sub>::mCherry; p<sub>hlh-3</sub>::TIR-1::mRuby]* | This study |
| Figure 4 |  |  |
| UD925 | *cgef-1(yc3)*, *unc-84(n369)*, *oxIs12 X*; *ycEx60*; *ycEx285[100ng/uL odr-1::gfp; 2ng/uL pSL884]* | This study |
| UD926 | *cgef-1(yc3)*, *unc-84(n369)*, *oxIs12 X*; *ycEx60*; *ycEx286[100ng/uL odr-1::gfp; 2ng/uL pSL884]* | This study |
| UD927 | *cgef-1(yc3)*, *unc-84(n369)*, *oxIs12 X*; *ycEx60*; *ycEx287[100ng/uL odr-1::gfp; 2ng/uL pSL884]* | This study |
| UD932 | *cgef-1(yc3)*, *unc-84(n369)*, *oxIs12 X*; *ycEx60*; *ycEx288[100ng/uL odr-1::gfp; 2ng/uL pSL885]* | This study |
| UD933 | *cgef-1(yc3)*, *unc-84(n369)*, *oxIs12 X*; *ycEx60*; *ycEx289[100ng/uL odr-1::gfp; 2ng/uL pSL885]* | This study |
| UD934 | *cgef-1(yc3)*, *unc-84(n369)*, *oxIs12 X*; *ycEx60*; *ycEx290[100ng/uL odr-1::gfp; 2ng/uL pSL885]* | This study |
| UD935 | *cgef-1(yc3)*, *unc-84(n369)*, *oxIs12 X*; *ycEx60*; *ycEx291[100ng/uL odr-1::gfp; 2ng/uL pSL886]* | This study |
| UD936 | *cgef-1(yc3)*, *unc-84(n369)*, *oxIs12 X*; *ycEx60*; *ycEx292[100ng/uL odr-1::gfp; 2ng/uL pSL886]* | This study |
| UD937 | *cgef-1(yc3)*, *unc-84(n369)*, *oxIs12 X*; *ycEx60*; *ycEx293[100ng/uL odr-1::gfp; 2ng/uL pSL886]* | This study |
| UD938 | *cgef-1(yc3)*, *unc-84(n369)*, *oxIs12 X*; *ycEx60*; *ycEx294[100ng/uL odr-1::gfp; 2ng/uL pSL887]* | This study |
| UD939 | *cgef-1(yc3), unc-84(n369), oxIs12 X; ycEx60; ycEx295[100ng/uL odr-1::gfp; 2ng/uL pSL887]* | This study |
| UD940 | *cgef-1(yc3), unc-84(n369), oxIs12 X; ycEx60; ycEx296[100ng/uL odr-1::gfp; 2ng/uL pSL887]* | This study |
| Figure 5 |  |  |
| BOX409 | *par-6(mib30[par-6::degron::egfp-loxp]) I; mibIs49[p<sub>wrt-2</sub>::TIR-1::tagBFP2-Lox511::tbb-2-3'UTR, IV:5014740-5014802 (cxTi10882 site)]) IV* | (Castiglioni et al., 2020) |
| BOX607 | *pkc-3(mib78[egfp-loxp::degron::pkc-3]) II; mibIs49[p<sub>wrt-2</sub>::TIR-1::tagBFP2-Lox511::tbb-2-3'UTR, IV:5014740-5014802(cxTi10882 site)]) IV* | (Castiglioni et al., 2020) |
| UD829 | *par-6(mib30[par-6::degron::egfp-loxp]) I; oxIs12 X; ycEx266[p<sub>myo-2</sub>::mCherry; p<sub>hlh-3</sub>::TIR-1::mRuby]* | This study |
| UD830 | *par-6(mib30[par-6::degron::egfp-loxp]) I; unc-84(n369), oxIs12 X; ycEx266[p<sub>myo-2</sub>::mCherry; p<sub>hlh-3</sub>::TIR-1::mRuby]* | This study |
| UD831 | *pkc-3(mib78[egfp-loxp::aid::pkc-3]) II; oxIs12 X; ycEx266[p<sub>myo-2</sub>::mCherry; p<sub>hlh-3</sub>::TIR-1::mRuby]* | This study |
| UD832 | *pkc-3(mib78[egfp-loxp::aid::pkc-3]) II; unc-84(n369), oxIs12 X; ycEx266[p<sub>myo-2</sub>::mCherry; p<sub>hlh-3</sub>::TIR-1::mRuby]* | This study |
| Figure 6 |  |  |
| DLW29 | *dpy-10, wLZ32[p<sub>sun-1</sub>::TIR-1::mRuby, Cbr-unc-119(+)]; arx-3(lib7[degron::arx-3] III, unc-119(ed3))* | (Zhang et al., 2015) |
| UD709 | *arx-3(lib7[degron::arx-3]) III; oxIs12 X; ycEx253[odr-1::rfp; p<sub>hlh-3</sub>::TIR-1::mRuby]* | This study |
| UD710 | *arx-3(lib7[degron::arx-3]) III; unc-84(n369), oxIs12 X; ycEx253[odr-1::rfp; p<sub>hlh-3</sub>::TIR-1::mRuby]* | This study |
| UD825 | *nmy-2(yc111[degron::GFP11::nmy-2]) I; oxIs12 X; ycEx266[p<sub>myo-2</sub>::mCherry; p<sub>hlh-3</sub>::TIR-1::mRuby]* | This study |
| UD826 | *nmy-2(yc111[degron::GFP11::nmy-2]) I; unc-84(n369), oxIs12 X; ycEx266[p<sub>myo-2</sub>::mCherry; p<sub>hlh-3</sub>::TIR-1::mRuby]* | This study |

For the cgef-1 RNAi experiments, clone X-2A03 from the Ahringer RNAi library (Source Bioscience) (Kamath et al., 2003) was used to create dsRNA *in vitro*, which was then injected into UD87 as described (Chang et al., 2013; Fire et al., 1998).

Fosmids used in the *cgef-1* rescue experiments were from the *C. elegans* Fosmid Library (Source BioScience) and were amplified in bacteria using CopyControl Induction Solution (Lucigen, #CCIS125) and purified using a DNA midi prep kit (Thermo Scientific, #K0481). Fosmid injection mixes contained 5 ng/μl of each indicated fosmid and 100 ng/μl of *odr-1::gfp* plasmid (L’Etoile & Bargmann, 2000) and were injected into UD285.

### CRISPR/Cas9 gene editing

*cgef-1* isoform mutants were generated using *dpy-10* as a co-CRISPR marker (Arribere et al., 2014; Paix et al., 2015, 2017). The CRISPR injection mix was generated as described (Hao et al., 2021). The same guides were used to create the deletion mutations of each isoforms but without the addition of the repair templates. Deletion mutations were screened by amplifying the region around the guide and PCR products that showed a smaller band size were sent for Sanger sequencing.

All crRNA and ssODN repair templates are listed in Table 2. Degron and GFP11 insertions for *cdc-42* and *nmy-2* were generated by using *zen-4(+)* as a co-CRISPR marker (Farboud et al., 2019). Single stranded repair templates for insertions contained 50nt homology arms (Genewiz). The CRISPR injection mix contained 0.084 μl *zen-4* crRNA (0.6mM), 0.21 μl target gene crRNA (0.6mM), 1.033 μl tracr (0.17mM), 4.39 μl Cas9 (40μM), 0.28 μl ssODN *zen-4(+)* repair template (500ng/μl), and 4 μl ssDNA repair template (500 ng/μl). The injection mix was injected into germline of temperature sensitive *zen-4(cle10)* mutant young adults and screened as previously described (Farboud et al., 2019).

**Table 2.**
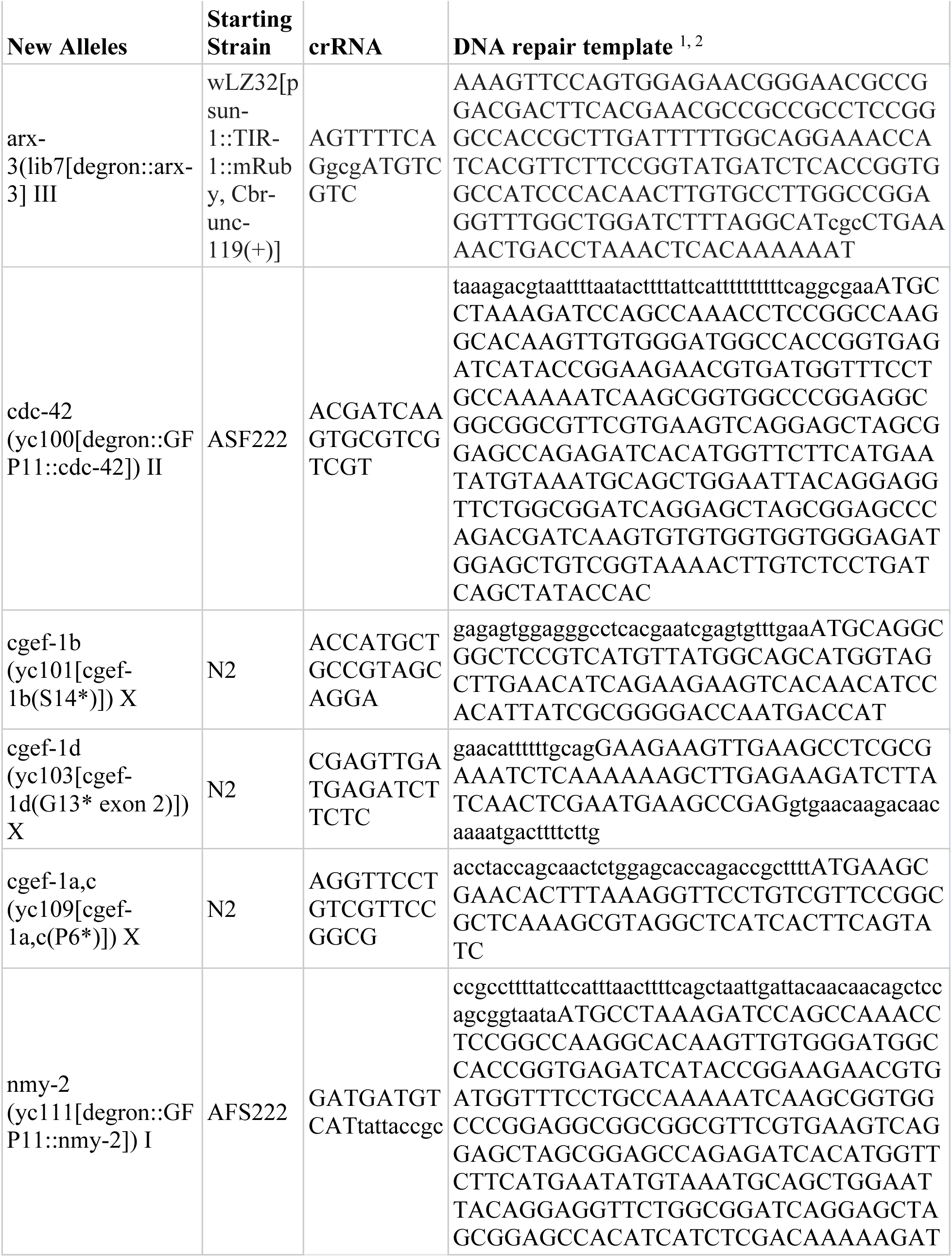
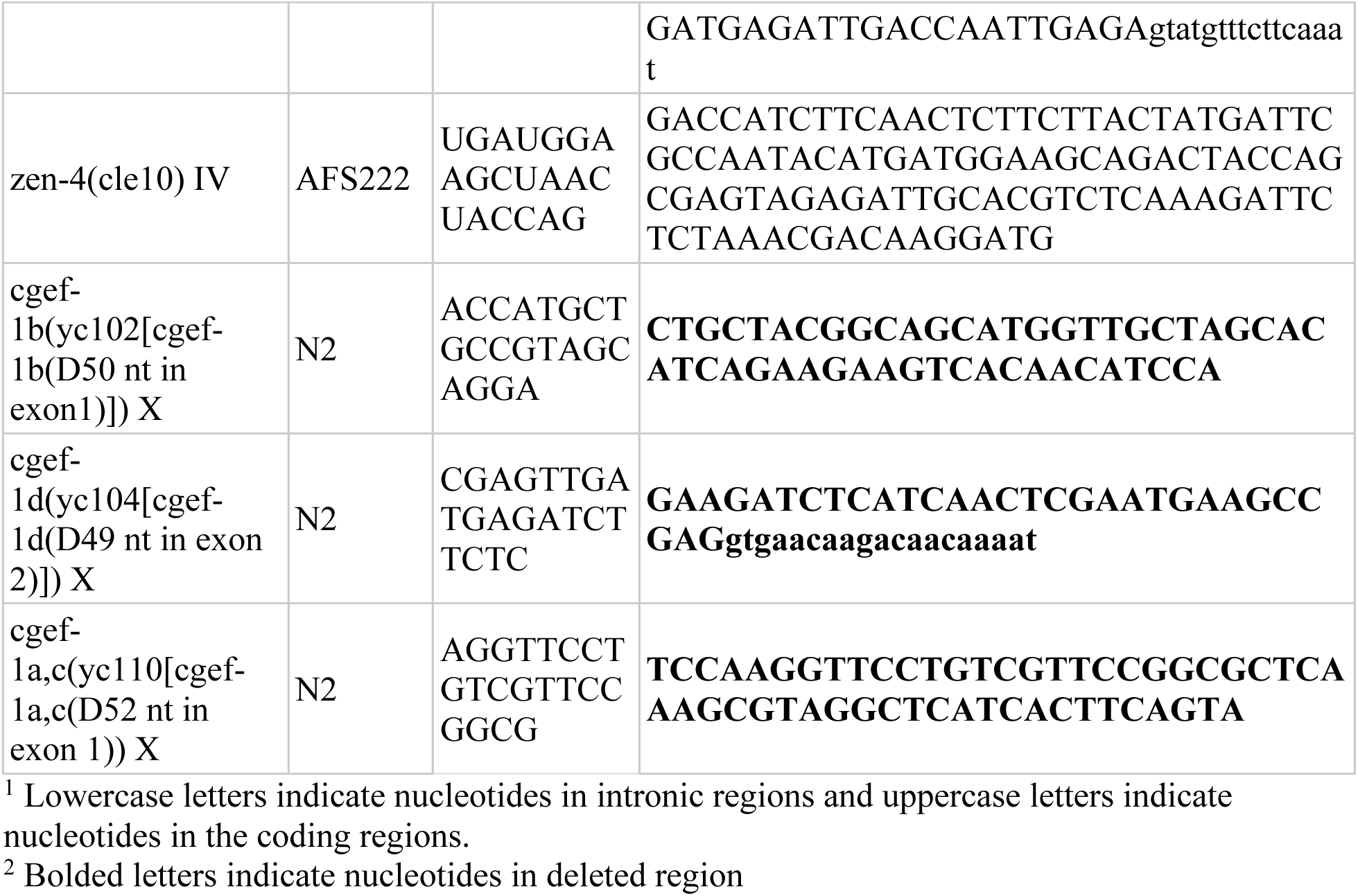
(CRISPR cRNA/Repair Templates)

An auxin-induced degron was inserted to the 5’ end of *arx-3*, replacing the native start codon of the gene. No linker sequence was used. The tag was inserted with a ssDNA oligo (Table 2). A *dpy-10* co-CRISPR strategy was used to identify successful injections and CRISPR repair activity (Arribere et al., 2014). The strain wLZ32[*p_sun-1_::TIR-1::mRuby, Cbr-unc-119(+)*] an unnamed strain from Abby Dernburg that expresses single copy TIR1 in the germline, was used for injections. Protein Cas9 and synthetic RNA was generated by Integrated DNA Technologies.

### Cloning constitutively active small Rho GTPase constructs

To generate plasmid pSL884 (*p_hlh-3_::2xHA::cdc-42(G12V)::unc-54 3’UTR*), the *cdc-42(G12V)* open reading frame was amplified from pEL298 (Alan et al., 2013) with homology arms to add a 2xHA tag after the *cdc-42* start codon. To generate plasmid pSL885 (*p_hlh-3_::2xHA::mig-2(G16V)::unc-54 3’UTR*), the *mig-2(G16V)* open reading frame was amplified from pEL656 (Norris et al., 2014) with homology arms to add a 2xHA tag after the *mig-2* start codon. To generate plasmid pSL886 (*p_hlh-3_::2xHA::ced-10(G12V)::unc-54 3’UTR*), the *ced-10(G12V)* open reading frame was amplified from pEL777 (Norris et al., 2014) with homology arms to add a 2xHA tag after the *ced-10* start codon. To generate plasmid pSL887 (*p_hlh-3_::2xHA::rho-1(G14V)::unc-54 3’UTR*), the *rho-1(G14V)* open reading frame was amplified from pEL1021 (Gujar et al., 2019) with homology arms to add a 2xHA tag after the *rho-1* start codon. The backbone of pSL830 including the promoter of *hlh-3* and the *unc-54 3’UTR* was amplified, and the HiFi DNA Assembly Cloning Kit (New England Biolabs) was used to assemble pSL884, pSL885, pSL886, and pSL887. Injection mixes containing 2 ng/μl of a plasmid encoding a constitutively active construct and 100 ng/μl of plasmid *odr-1::gfp* (L’Etoile & Bargmann, 2000) as a co-injection marker were injected into UD285. A list of plasmids for transgenic constructs are listed in Table 3.

**Table 3.** (Reagents)

| Reagent Type | Name | Source or reference | Additional Information |
| --- | --- | --- | --- |
| *Escherichia. coli* | OP50 | Caenorhabditis Genetics Center (CGC) |  |
| Recombinant DNA reagent | pSL619 | (Chang et al., 2013) | p<sub>hlh-3</sub>::nls::tdTOMATO |
| Recombinant DNA reagent | pSL814 | (J. Ho et al., 2018) | p<sub>hlh-3</sub>::TIR-1::mRuby |
| Recombinant DNA reagent | pSL830 | This paper | p<sub>hlh-3</sub>::GFP(1-10) |
| Recombinant DNA reagent | pSL884 | This paper | p<sub>hlh-3</sub>::2xHA::cdc-42(G12V) |
| Recombinant DNA reagent | pSL885 | This paper | p<sub>hlh-3</sub>::2xHA::mig-2(G16V) |
| Recombinant DNA reagent | pSL886 | This paper | p<sub>hlh-3</sub>::2xHA::ced-10(G12V) |
| Recombinant DNA reagent | pSL887 | This paper | p<sub>hlh-3</sub>::2xHA::rho-1(G14V) |
| Recombinant DNA reagent | pLZ31 | (Zhang et al., 2015) | p<sub>eft-3</sub>::TIR-1::mRuby |
| Recombinant DNA reagent | pEL298 | (Alan et al., 2013) | p<sub>osm-6</sub>::cdc-42(G12V)::GFP |
| Recombinant DNA reagent | pEL656 | (Norris et al., 2014) | p<sub>unc-25</sub>::mig-2(G16V)::GFP |
| Recombinant DNA reagent | pEL777 | (Norris et al., 2014) | p<sub>unc-25</sub>::ced-10(G12V)::GFP |
| Recombinant DNA reagent | pEL1021 | (Gujar et al., 2019) | p<sub>unc-25</sub>::rho-1(G14V)::GFP |

### P-cell nuclear migration assay

For the P-cell nuclear migration assays, *oxIs12[p_unc-47_::gfp]* transgenic worms (EG1285) were used as the wild type control (McIntire et al., 1997). *oxIs12* was used as a reporter for P-cell derived GABA neurons to assay for P-cell nuclear migration defects. L4 animals were mounted onto 2% agarose slides in 1mM tetramisole solution. Slides were viewed using a wide-field epifluorescent Leica DM6000 microscope and a 63x Plan Apo 1.40 NA objective. UNC-47::GFP-positive GABA neurons were counted in the ventral cord. Neurons normally outside the ventral cord in the nerve ring and the most posterior neuron in the tail are not decedents of P cells and were not counted. A total of 19 GABA neurons (12 derived from P cells) were scored for each animal as previously described (Fridolfsson et al., 2018).

### Synchronization and auxin assay

We synchronized *C. elegans* larvae in the mid L1 stage at approximately the time of P-cell nuclear migration for the auxin experiments. 50-100 L4s were picked and grown at 20°C for 24-48 hours so that animals reached the adult stage. Adult animals were transferred onto a fresh NGM plate and allowed to lay eggs at 20°C for 1 hour. After 1 hour, the adult animals were removed, leaving synchronized embryos behind.

Conditional knock-down of proteins of interest was achieved using the auxin-inducible degradation system (Zhang et al., 2015). TIR-1 was amplified from plasmid pLZ31 (Zhang et al., 2015) (Addgene #71720) and cloned under control of the *hlh-3* promoter in pSL780 (Bone et al., 2016) with Gibson cloning (New England Biolabs) to generate pSL814. pSL814 was injected with *odr-1::rfp* to make the extrachromosomal arrays in strains UD709 and UD710. pSL814 was injected with *p_myo-2_::mCherry* to make the extrachromosomal arrays in strains UD716, UD717, UD825, UD826.

Synchronized L1 animals were washed off normal NGM plates with distilled water approximately two hours before P-cell nuclear migration began. Next, the L1 larvae were transferred to NGM+auxin plates with 1 mM 3-indoleacetic acid (IAA; Sigma #I2886) and kept in the dark. After P-cell nuclear migration was completed, the L1 larvae were washed off the NGM+auxin plates with M9 buffer and subsequently transferred to an NGM plate. For experiments performed at 15°C, embryos were left on the plates to develop for 29 hours and then the resulting L1s were washed onto NGM + auxin plates and left to develop for 8 hours before being washed off onto NGM plates without auxin to develop to the L4 stage. For experiments done at 20°C, embryos developed into L1s on NGM plates for 18 hours and then placed on NGM with auxin plates for 7 hours. For experiments performed at 25°C, embryos developed into L1s for 12 hours and then placed on NGM with auxin plates for 6 hours. These timings were determined by using the marker p_hlh-3_::nls::tdTOMATO (pSL619) to visualize P-cell nuclei throughout development. Once animals reached the L4 stage, the number of UNC-47::GFP marked GABA neurons were quantified.

To synchronize *C. elegans* for imaging, 3-5 plates of gravid animals were bleached and eggs were pelleted and resuspended in M9 solution for 12-16 hours at room temperature on a rocker. After starvation, L1s were plated onto NGM plates seeded with OP50 and grown at room temperature for 12 hours. After 12 hours, L1s were washed off the plates with M9 and mounted onto 2% agarose slides with 1mM tetramisole solution.

### Microscopy Imaging

Images were taken on a Zeiss LSM 980 with Airyscan using 20x objective and the Zeiss Zen Blue software which was provided by the MCB light imaging microscopy core and by NIH grant S10OD026702.

### Statistical Analysis

Graphs of GABA neuron counts were created by using Prism (version 9). An unpaired, two-tailed student t-test was used to determine statistical significance. Error bars are 95% confidence intervals.

## Acknowledgements

We thank past and present members of the Starr and Luxton lab for their input on this research. We thank Zach Stevenson for help making a strain. We thank David Sherwood, Mike Boxem, Abby Dernburg, and the Caenorhabditis Genetics Center (CGC), which is funded by the National Institutes of Health Office of Research Infrastructure Programs (P40OD010440), for providing strains. We thank WormBase. This research was funded by a grant from the National Institutes of Health, R35GM134859 to D.A.S. and R35GM128890 to D.L.

